# Endogenous oxytocin inhibits hypothalamic stress responsive neurons following acute hypernatremia

**DOI:** 10.1101/543165

**Authors:** Dipanwita Pati, Scott W. Harden, Kyle B. Kelly, Annette D. de Kloet, Eric G. Krause, Charles J. Frazier

## Abstract

Significant prior evidence indicates that centrally acting oxytocin robustly modulates stress responsiveness and anxiety-like behavior, although the neural mechanisms behind these effects are not completely understood. A plausible neural basis for oxytocin mediated stress reduction is via inhibition of corticotropin-releasing hormone (CRH) neurons in the paraventricular nucleus of the hypothalamus (PVN) that regulate activation of the hypothalamic-pituitary-adrenal (HPA) axis. Previously, we have shown that following subcutaneous injection of 2.0 M NaCl, oxytocin (OT) synthesizing neurons are activated in the rat PVN, an oxytocin receptor (Oxtr) dependent inhibitory tone develops on a subset of parvocellular neurons, and stress-mediated increases in plasma corticosterone levels are blunted. Here, we utilized transgenic male CRH-reporter mice to selectively target PVN CRH neurons for whole-cell recordings. These experiments reveal that acute salt loading produces tonic inhibition of PVN CRH neurons through a mechanism that is largely independent of synaptic activity. Further studies reveal that CRH neurons within the PVN synthesize mRNA for Oxtr(s). Salt induced Oxtr-dependent inhibitory tone was eliminated in individual PVN CRH neurons filled with GDP-β-S, and was also largely absent in PVN CRH neurons extracted form CRH-Oxtr KO mice. Additional electrophysiological studies suggest that reduced excitability of PVN CRH neurons in salt loaded animals is associated with increased activation of an inwardly rectifying potassium channel. Collectively, these data reveal a likely cellular mechanism by which endogenous oxytocin signaling reduces the excitability of PVN CRH neurons to curb stress responsiveness during times of high plasma osmolality.

## Introduction

The neuroendocrine response to acute stress is largely defined by changes in the state of activation of the sympathetic nervous system and of the hypothalamic-pituitary-adrenal (HPA) axis. Both effects depend in large part on modulation of distinct populations of neurons within the paraventricular nucleus of the hypothalamus (PVN). These include preautonomic parvocellular neurons, which project to preganglionic sympathetic neurons in the hindbrain and spinal cord to regulate sympathetic output, and corticotropin releasing hormone (CRH) positive parvocellular neurosecretory neurons, which project to the median eminence to regulate activation of the HPA axis. Both populations of neurons are subject to a modulatory control by a wide range of endogenous and exogenous compounds.

Oxytocin is one such neuromodulator, an endogenous neuropeptide produced predominantly in the paraventricular and supraoptic nuclei of the hypothalamus. Significant prior evidence suggests that endogenous oxytocin, acting in the PVN, is likely to directly modulate stress induced activation of the HPA axis. Indeed, a variety of distinct psychological stressors associated with increased HPA activity promote central release of oxytocin (Ježová et al., 1993; Nishioka et al., 1998; Wotjak et al., 1998; Engelmann et al., 1999; Torner et al., 2017), chronic administration of oxytocin into the PVN blunts stress responsiveness (Windle et al., 1997, 2004), and intracerebral infusion of an Oxtr antagonist promotes stress induced release of both adrenocorticotropic hormone (ACTH) and corticosterone into the blood (Neumann et al., 2000). Similarly, oxytocin knockout mice produce elevated levels of CRH mRNA in the PVN in response to restraint stress (Nomura et al., 2003), while lactating rodents demonstrate both elevated levels of oxytocin release and reduced levels of CRH mRNA in the PVN (Bealer and Crowley, 1998; Walker et al., 2001). Collectively, these data suggest that oxytocin release, produced in response to psychological stressors, initiates a negative feedback mechanism that ultimately dampens stress responsiveness in part through down regulation of CRH synthesis in the PVN (Jurek et al., 2015; Winter and Jurek, 2018).

Intriguingly, several findings from our group suggest that oxytocin release, produced by a stressor that is not associated with robust activation of the HPA axis, likely also dampens stress responsiveness, but through a mechanism not likely to involve genomic modulation of CRH synthesis. Specifically, we have reported that acute hypernatremia, a peripheral stressor that promotes vasopressin dependent antidiuresis and oxytocin dependent natriuresis (Huang et al., 1996; öquist et al., 1999; McKinley et al., 2004; Tucker and Stocker, 2016), also directly reduces HPA axis activation produced by subsequent restraint stress (Krause et al., 2011). Notably, additional work in both rats and mice revealed that this effect is associated with increased activation of c-Fos in PVN oxytocinergic neurons, decreased activation of c-Fos in PVN CRH neurons, and development of an Oxtr dependent inhibitory tone on a subset of PVN parvocellular neurons that lack both a prominent A-type potassium current and a robust low threshold spike (Frazier et al., 2013; Smith et al., 2014, 2015).

The intrinsic electrophysiological phenotype of the PVN neurons that were inhibited by oxytocin after acute hypernatremia in Frazier et al (2013) were consistent with the interpretation that they were CRH expressing neurosecretory neurons involved in regulation of the HPA axis (Luther et al., 2002), but this phenotype was not directly confirmed, and the mechanism of putative Oxtr mediated inhibition was not investigated. Therefore, in the current study we used male mice that express tdTomato in CRH neurons to directly test the hypothesis that acute hypernatremia creates an Oxtr-mediated inhibitory tone on PVN CRH neurons, and to further explore the cellular mechanisms and downstream effects of this inhibition. Our results indicate that Oxtr-dependent inhibitory tone is produced by acute hypernatremia in PVN CRH neurons, that the effect depends largely on Oxtr(s) specifically expressed by these neurons, that the Oxtr(s) in question are likely coupled to a potassium permeant ionophore, and that activation of this channel reduces neuronal gain.

## Materials and Methods

### Animals

#### CRH -reporter mice

Most studies were performed on CRH-reporter mice that expressed red fluorescent protein (tdTomato) in neurons that produce CRH. These mice were generated by crossing *Crh-IRES-Cre* mice (B6(Cb)-Crh^tm1(cre)Zjf^/J mice, Jackson Laboratories Stock #012704) with mice that express tdTomato behind a floxed stop codon in the Rosa26 locus under control of the CAG promoter (B6.Cg-Gt(ROSA)26Sor^tm14(CAG-TdTomato)Hze^/J, Jackson Laboratories Stock #007914). These same CRH-reporter mice have been previously found, by our laboratory and others, to express mRNA for CRH in 80-95% of tdTomato positive neurons in the hypothalamus (Cusulin et al., 2013; Smith et al., 2014).

#### CRH-Oxtr KO mice

Mice used in these experiments were on a C57BL/6J background. In order to generate mice that lack Oxtr(s) in CRH neurons and their littermate controls, mice homozygous for the Oxtr flox knock-in gene and also heterozygous for the CRH-Cre knock-in gene were bred with those expressing only the Oxtr-flox gene. Oxtr flox knock-in mice (Jackson laboratories; Stock # 008471) express loxP sites that flank exons 2-3 of the Oxtr gene (Lee et al., 2008). Additional details of these mouse lines are available at the vendor’s website.

### Saline injections and slice preparation

On the day of experimentation, male mice were assigned to one of two groups: isotonic or hypertonic. Mice in the isotonic group received subcutaneous injections of 0.1 mL of 0.15 M NaCl, while mice in the hypertonic group received subcutaneous injections of 0.1 mL of 2.0 M NaCl. All animals were euthanatized 1-hour post injection. Mice in the isotonic group were allowed free access to water in the one hour between injection and euthanasia, while mice in the hypertonic group were denied access to water between the time of injection and euthanasia. To minimize pain and irritation, each injection was preceded by 2% lidocaine (∼0.01 mL). Prior work from our group has revealed that this protocol increases plasma sodium concentration without affecting indices of blood volume (Smith et al., 2014). Sixty minutes after saline injections, mice were administered ketamine (80–100 mg/kg, ip) and were rapidly decapitated using a rodent guillotine. The brain was quickly removed, and coronal sections (300 μm thick) through the PVN were made using a Leica VT 1000s vibratome. Slices were incubated for 30 minutes in a dissecting solution maintained at 30-35°C that contained in mM: 124 NaCl, 2.5 KCl, 1.23 NaH2PO4, 2.5 MgSO4, 10 D-glucose, 1 CaCl2, and 25.9 NaHCO3, saturated with 95% O2-5% CO2. After equilibrating at room temperature for at least 30 minutes, slices were transferred to a slice chamber for experimental use.

### *In Vitro* Electrophysiology

For whole-cell recording, slices were continuously perfused at a rate of 1.2-1.5 mL/min with aCSF that contained (in mM): 126 NaCl, 3 KCl, 1.2 NaH_2_PO_4_, 1.5 MgSO4, 11 D-glucose, 2.4 CaCl_2_, and 25.9 NaHCO_3_. This solution was saturated with 95% O_2_ and 5% CO_2_, and bath temperature was maintained at 28±2°C. For experiments that required minimal spontaneous synaptic activity in the slice, this ACSF was supplemented with TTX (1µM), to block voltage-gated sodium channels, DNQX (20 µM) and DL-2-amino-5-phosphonopenatanoic acid (APV, 40 µM), to block ionotropic glutamate receptors, picrotoxin (100 µM) to block both synaptic and extrasynaptic GABA_A_ receptors. For most experiments the patch electrode was filled with a K-gluconate based internal solution that contained (in mM): 130 K-gluconate, 10 KCl, 10 NaCl, 2 MgCl_2_, 1 EGTA, 2 Na_2_ATP, 0.3 NaGTP, and 10 HEPES, pH adjusted to 7.3 using KOH and volume adjusted to 285–300 mOsm. For experiments that involved intracellular inhibition of G-protein coupled receptors, the GTP in this internal was replaced with GDP-β-S (300 µM), and cells were allowed to equilibrate for 30 minutes after whole-cell configuration was attained to allow intracellular diffusion of GDP-β-S into the cell from the patch pipette. For experiments designed to measure spontaneous synaptic currents, a Cs-gluconate based internal solution was used, which contained, in mM: 140 Cs-gluconate, 3 NaCl, 1 MgCl_2_, 0.2 EGTA, 4 Na_2_ATP, 0.3 NaGTP, 5 QX-314-Cl, and 10 HEPES, pH adjusted to 7.3 using CsOH and volume adjusted to 285–300 mOsm. In all cases, slices were visualized using an Olympus BX51W1 microscope using a combination infrared differential interference contrast (IR DIC) and epifluorescence microscopy. Whole-cell voltage-clamp recordings were performed using micropipettes pulled from a borosilicate glass using a Flaming/Brown electrode puller (Sutter P-97; Sutter Instruments, Novato, California). Electrode open tip resistance was between 4 and 6 MΩ using the K-gluconate internal. Voltage-clamp experiments were performed using an Axon Multiclamp 700A or 700B amplifier (Molecular Devices, Sunnyvale, CA). Data were sampled at 20 kHz, filtered at 2 kHz, and recorded on a computer by a Digidata 1400 A/D converter using Clampex version 10 (Molecular Devices, Sunnyvale, CA). All chemicals used for in vitro electrophysiology were obtained from either Tocris Cookson or Sigma-Aldrich except [d(CH2)51,Tyr(Me)2,Orn8]-oxytocin (Oxtr-A), which was obtained from Bachem.

### Immunohistochemistry and two-photon microscopy

Immunohistochemical studies were performed on 300 µm brain slices prepared identically to those used for *in-vitro* electrophysiology. Slices were preserved in 10% formalin overnight at 4C. All subsequent steps were performed at 25C on an orbital shaker. Slices were washed with PBST (PBS + 0.1% Triton X-100) 5x 5 min, then incubated in blocking solution (PBST supplemented with 2% bovine serum albumin and 2% normal goat serum) for 48hr. Primary antibodies against NP1 (1:500 goat anti-mouse neurophysin-1, Millipore MABN844) and tdTomato (1:500 rabbit anti-RFP, Rockland 6004013795) were then applied in blocking solution for 72hr. Slices were washed (5x 5 min with PBST) then incubated in secondary antibodies against mouse (1:500 goat anti-mouse, ThermoFisher A11001) and rabbit (1:500 goat anti-rabbit, ThermoFisher A11012) conjugated with two-photon-excitable fluorophores (488nm and 594nm, respectively) for 24hr. Slices were washed (5x 5 min with PBS), coverslipped with an aqueous mounting medium (Vectashield) and imaged with a two-photon laser scanning microscope (excited by 810 nm femtosecond laser).

### RNAscope in situ hybridization

RNAscope in situ hybridization (Advanced Cell Diagnostics, Newark, CA, USA) was performed on brain sections of CRH-tdTomato mice as per the manufacturer’s instructions, using the same modifications that we have previously described (Smith et al., 2014; de Kloet et al., 2016). These modifications allow for the visualization of mRNA transcripts (visible as punctate dots) in the presence of preserved tdTomato fluorescence. Briefly, mice were euthanized with an overdose of sodium pentobarbital and transcardially perfused with 4% RNase free paraformaldehyde (PFA). Brains were extracted, post-fixed in 4% PFA for 4 h, and then transferred into RNase free 30% sucrose at 4° C. Brains were then coronally sectioned at 20 µm using a cryostat. Brain sections were rinsed twice in RNase free PBS, mounted onto Superfrost Plus Gold slides, air-dried for 20 min at room temperature and then stored at −80° C. On the day of RNAscope in situ hybridization, brain sections were again air-dried for 30 min at room temperature, incubated in protease IV for 20 min, and then hybridized with specific probes. Probes used in this study were: (1) DapB, negative control, (2) Ubc, positive control, and (3) Oxtr. The probe for Oxtr was used to determine the percentage of CRH-tdTomato neurons in the PVN that express Oxtr. The color label for all probes was assigned to FAR RED (Excitation: 647 nm; Emission: 690 ± 10 nm).

### Image capture, processing and analysis

For RNAscope in situ hybridization studies, z-stacks throughout the PVN were captured and processed using Axiovision 4.8.2 software and a Zeiss AxioImager fluorescent Apotome microscope (Carl Zeiss Microscopy, Thornwood, NY, USA). In all cases, z-stacks of tdTomato fluorescence and transcripts of interest were captured at 40× magnification. These z-stacks each contained an average of 20 optical sections, with the distance between the optical sections set at 0.5 µm. For each mouse, an average of 4 z-stacks were captured throughout the rostral-caudal extent of the parvocellular neurosecretory region of the PVN. Importantly, when capturing these images, sections hybridized with the positive control probes were used to determine the exposure time and image processing required to provide optimal visualization of RNA signal. Then, as described in detail previously (de Kloet et al., 2016), these same parameters were used for visualization of mRNA transcripts of interest and to assess background fluorescence in sections hybridized with negative control probe (DapB). Importantly, using these exposure times and image processing parameters there was minimal or no fluorescence in sections hybridized with the negative control probe. The projection image shown in the results section was generated from a representative z-stack and then prepared using Adobe Photoshop 7.0 where the brightness and contrast was adjusted to provide optimal visualization.

Analysis of co-localization of Oxtr mRNA transcripts with tdTomato fluorescing cells was performed manually on the 40 × magnification z-stacks through the parvocellular PVN of 3 separate CRH-tdTomato mouse brains. TdTomato neurons were considered to contain the RNA of interest if at least 2 visible transcripts, defined as an individual punctate dot, were observed within the volume of the tdTomato fluorescence. Data are then reported as the percentage of tdTomato cells that contain the Oxtr mRNA for Oxtr within the parvocellular PVN.

### Data and statistical analysis

The analysis of all electrophysiological data was performed using custom software written in OriginC (OriginLab, Northampton, MA) by CJF. Current density was calculated as the ratio of holding current to whole cell capacitance (pA/pF) when cells were continuously voltage clamped at −70 mV. Spontaneous postsynaptic currents were detected using parameter based event detection software. Event parameters were chosen to detect the vast majority of distinguishable events while minimizing false positives. Typical threshold amplitude was 6-8 pA and typical threshold area was 20-40 pA*ms. In all cases an algorithm for detecting complex peaks (peaks that occur during the decay period of a previous event) was employed. This algorithm calculates event amplitude for the second (and subsequent) peaks within the decay period based on extrapolated monoexponential decay from the peak of the initial event. Within groups, a two-tailed, one-sample t-test was used to evaluate changes in current density following the antagonism of oxytocin receptors [null hypothesis, Δ current density = 0]. In experiments that involved comparing a single variable across two groups, two-tailed, two-sample unpaired t-test was used, and Welch’s correction was applied in cases where the groups had unequal variance. For statistical analyses in Fig. 6A-B and E-G, a two-way repeated measures ANOVA was employed (salt condition [isotonic or hypertonic] × current or voltage command). When this analysis revealed a main effect of salt loading and/or a significant salt × command interaction, Sidak-Holm post-hoc tests were used to compare individual means. For all electrophysiological experiments testing the effects of bath applied drugs, baseline was considered as 2 −5 mins after initiation of a stable recording, and the drug effect was measured either between 11-13 mins or 9-11 mins after onset of drug application. All data are expressed as mean ± SEM. P-values ≤ 0.05 were considered significant.

## Results

### Acute salt loading produces an oxytocin receptor dependent tonic inhibitory current in PVN CRH neurons

Prior work from our group revealed that acute hypernatremia reduces activation of the HPA axis in response to subsequent restraint stress, and produces an inhibitory oxytocinergic tone in a subset of PVN parvocellular neurons (Krause et al., 2011; Frazier et al., 2013; Smith et al., 2014). In order to determine whether this phenomenon involves direct inhibition of PVN CRH neurons, we crossed Crh-IRES-Cre mice with a tdTomato Cre reporter to create a CRH reporter animal (see methods). Prior work from our group and others has confirmed that tdTomato expression is an excellent marker of PVN CRH neurons in this animal model (Cusulin et al., 2013; Smith et al., 2014; Chen et al., 2015; Walker et al., 2018). Immunohistochemistry for neurophysin 1, an oxytocin transporter, in CRH reporter animals revealed that PVN CRH neurons are located in close proximity to NP1 positive and thus presumably oxytocinergic neurons, and also further reinforce that there is minimal overlap between these two populations (Fig. 1A). We performed whole-cell patch clamp recordings from PVN CRH neurons from these animals identified with a combination of epifluorescence and differential interference contrast microscopy (Fig. 1B-D). Intrinsic properties of CRH neurons were calculated from data generated in voltage clamp using brief voltage steps from −70 to −80 mV. Consistent with expectations, we found the vast majority of PVN CRH neurons had high input resistance (typically > 800 MΩ), small whole-cell capacitance (typically < 20 pF), lacked a robust transient outwardly rectifying potassium current (I_A_), typical of magnocellular neurons, and also lacked a robust low threshold spike, often present in preautonomic neurons (Luther et al., 2002; Cusulin et al., 2013; Frazier et al., 2013).

**Figure 1:**
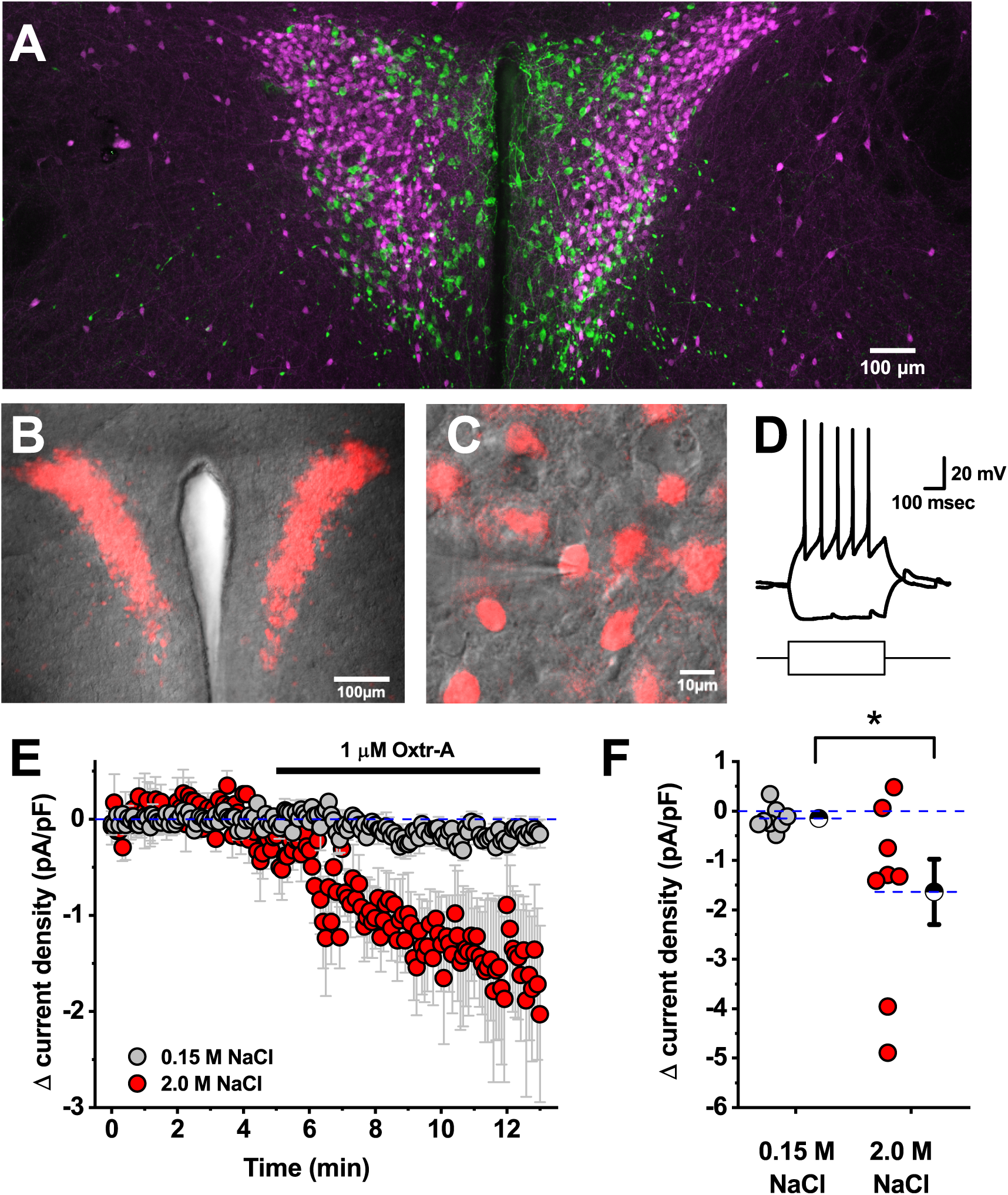
Acute salt loading produces an oxytocin receptor dependent tonic inhibitory current in PVN CRH neurons. **A)** Immunohistochemistry for NP1 in CRH reporter animals suggest that PVN oxytocinergic neurons (green) are located in close proximity to PVN CRH neurons (magenta), but that there is minimal if any overlap between these populations. **B)** tdTomato (red), as a marker for CRH neurons, can also be visualized in combination with DIC (gray) in an *in-vitro* slice preparation using a combination of epifluoresence and differential interference contrast microscopy. **C)** CRH neurons can be selectively targeted for patch-clamp analysis by their fluorescence. The patch pipette if visible coming in from the left side of the image. **D)** Representative response of a CRH neuron to +/-20 pA current steps. The shape of the depolarizing voltage trace is characteristic of CRH neurons (lacking both an I_A_ current and an LTS). A small hyperpolarizing step does not reveal robust hyperpolarization activated currents. **E)** Mean change in current density in response to Oxtr-A in CRH neurons from mice which received isotonic or hypertonic saline injection (grey vs. red, respectively). **F)** Specific change in current density produced by Oxtr-A for each cell that contributed to D. Hatched symbols with error bars represent mean and SE of each group. Asterisk indicates a significant difference between groups (p=0.04).

In order to evaluate the effect of acute salt loading (see methods) on identified PVN CRH neurons we voltage-clamped the neurons at −70 mV and bath applied an oxytocin receptor antagonist (Oxtr-A, 1 µM). In animals that received isotonic saline, Oxtr-A had no effect on current density (current per unit capacitance) required to hold the cells at −70 mV. However, in animals that received injections of hypertonic saline, bath application of Oxtr-A produced a clear decrease in current density (−1.6 ± 0.7 pA/pF, n=8, p=0.04 vs. isotonic condition, Fig. 1D-E). This result effectively reveals that salt acute loading produces an oxytocin receptor dependent tonic inhibitory current in genetically identified PVN CRH neurons.

### Acute salt loading does not alter spontaneous synaptic neurotransmission to the parvocellular CRH neurons

We next asked whether salt loading might also modulate excitability of PVN CRH neurons by altering the frequency or amplitude of spontaneous excitatory or inhibitory synaptic currents in the PVN. In order to evaluate those possibilities, we used a Cs-gluconate based internal solution (see methods) to isolate glutamatergic spontaneous excitatory postsynaptic currents (sEPSCs) in PVN CRH neurons voltage clamped at −60 mV, and also to isolate GABAergic spontaneous inhibitory postsynaptic currents (sIPSCs) in the same cells when voltage clamped at +10 mV (Fig. 2A-B). Overall, these experiments revealed that acute salt loading had no significant effect on either frequency or amplitude of sEPSCs or sIPSCs as observed in PVN CRH neurons (Fig. 2C-D). Specifically, in slices extracted from animals that received isotonic saline the frequency of sEPSCs was 2.9 ± 0.7 Hz (n=9) while the frequency of sIPSCs was 2.1 ± 0.3 Hz (n=13). In slices extracted from animals that received hypertonic saline theses value were 2.0 ± 0.5 Hz for sEPSC frequency and 2.7 ± 0.7 Hz, for sIPSC freqeuncy (n= 9, 13, and p=0.29 and 0.43, respectively). Mean sEPSC and sIPSC amplitudes were similarly unaffected (see Fig. 2D-E).

**Figure 2:**
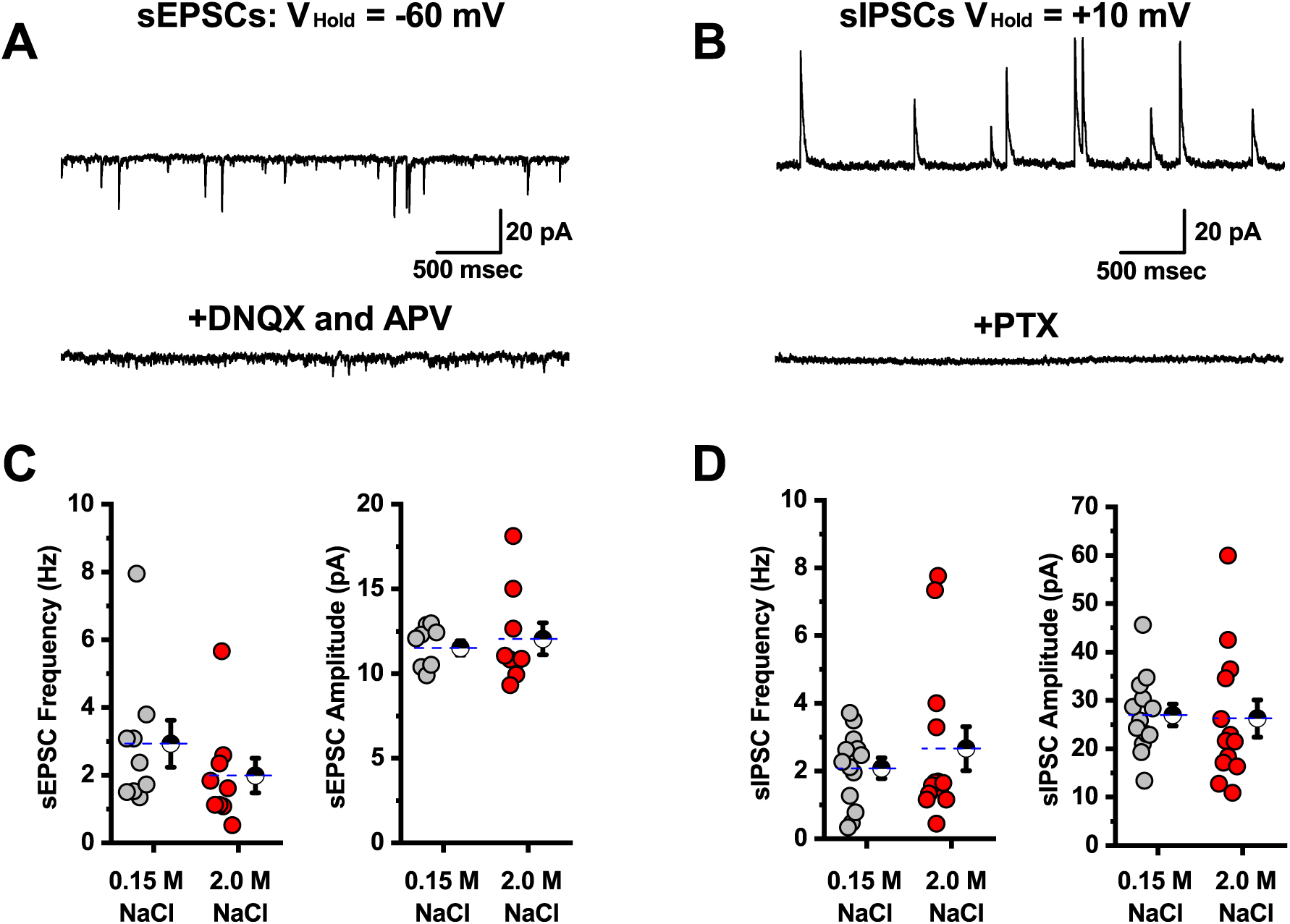
Acute salt loading does not alter spontaneous synaptic neurotransmission to PVN CRH neurons. **A)** Isolated sEPSCs in a representative PVN CRH neuron voltage clamped at −60 mV (top trace) are blocked by bath application of the ionotropic glutamate receptor antagonists DNQX and APV (bottom trace). **B)** Isolated sIPSCs in a representative PVN CRH neuron voltage clamped at +10 mV (top trace) are blocked by bath application of the GABA_A_ receptor antagonist PTX. **C)** Summary data illustrating sEPSC frequency and amplitude as observed on a cell by cell basis for cells voltage clamped at −60 mV. **D)** Summary data illustrating sIPSC frequency and amplitude as observed on a cell by cell basis for cells voltage clamped at +10 mV. Overall, there was no significant difference in the frequency or amplitude of spontaneous glutamatergic or GABAergic currents observed in PVN CRH neurons extracted from slices that received isotonic vs. hypertonic saline (grey vs. red, respectively). For each box plot, large hatched symbols to the right of the data represent the mean of each group, and error bars indicate the SE.

### Oxytocinergic receptor mediated tonic current in CRH neurons is insensitive to glutamate blockers, GABA blockers, and TTX

Although salt loading did not alter glutamate or GABA_A_ receptor mediated spontaneous synaptic transmission as observed in PVN CRH neurons, it seemed possible that oxytocin receptor activation could still indirectly modulate CRH neurons by altering transmitter uptake or breakdown in ways that influence activation of extrasynaptic receptors, or by promoting activity dependent release of a transmitter that does not act through glutamate or GABA_A_ receptors. In order to evaluate those possibilities, we repeated our experiments with Oxtr-A in slices extracted from animals that received hypertonic saline. Experiments were performed exactly as in Fig. 1D (red traces) except that we added DNQX, APV, and PTX, to block not only synaptic, but also extrasynaptic ionotropic glutamate and GABA_A_ receptors, and also TTX (1 μM) to block all activity dependent synaptic transmission in the slice. Under these conditions, bath application of Oxtr-A still produced a robust decrease in current density in PVN CRH neurons voltage clamped at −70 mV (−1.2 ± 0.3 pA/pF, n=5, p=0.02 on a one-sample t-test vs. null hypothesis of mean = 0, Fig. 3A-B). Notably, this effect is not significantly different than observed in Fig. 1D in the absence of DNQX, APV, PTX, and TTX (p=0.61, two-sample unpaired t-test). Collectively these results suggest that Oxtr-mediated tonic inhibition of PVN CRH neurons as observed in slices from salt loaded animals is independent of synaptic or extrasynaptic activation of ionotropic glutamate or GABA_A_ receptors and does not require ongoing action potential dependent synaptic transmission.

**Figure 3:**
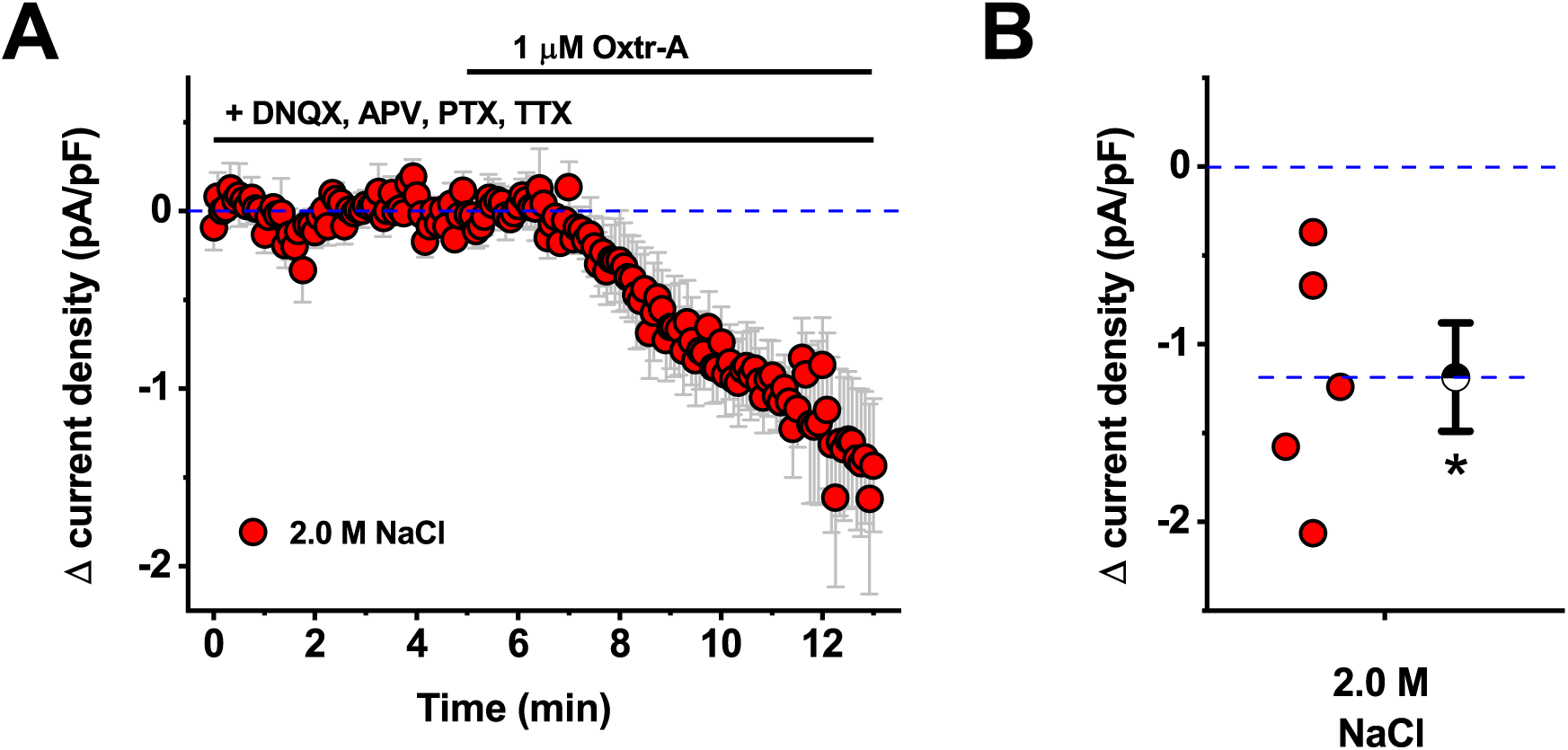
Oxytocin receptor mediated tonic current in CRH neurons is insensitive to glutamate blockers, GABA blockers, and TTX. **A)** Bath application of Oxtr-A continues to reveal an inhibitory tone in PVN CRH neurons extracted from animals that received hypertonic saline, even in the presence of bath applied antagonists for ionotropic glutamate receptors (DNQX and APV), GABA_A_ receptors (PTX), and voltage gated sodium channels (TTX). **B)** Box plot illustrating specific change in current density produced by Oxtr-A for each cell that contributed to the summary plot in panel A. Hatched symbol to the right of the data represents the population mean, and the error bars indicate the SE. The asterisk indicates mean change is significantly different than 0 (p=0.02).

### Inhibitory tone observed in CRH neurons from salt loaded animals is blocked by intracellular inhibition of GPCRs

Based on results presented above, we next sought to test the hypothesis that oxytocin receptor mediated tonic inhibition of PVN CRH neurons, as observed in slices extracted from salt loaded animals, is mediated by direct activation of G-protein coupled Oxtr(s) expressed by the PVN CRH neurons themselves. As a first step towards testing that hypothesis, we again repeated experiments in slices extracted from animals injected with hypertonic saline, as presented in Fig. 1D-E (red data plots), with two important changes. First, we used a more potent and selective oxytocin receptor antagonist, L-368,899-hydrochloride (L-368, 1 μM) in place of Oxtr-A (Busnelli et al., 2013). Second, in a subset of experiments we replaced the GTP in the internal solution with 300 μM GDP-β-S, a non-hydrolysable analog of GDP that competitively inhibits G-protein activation. As such, in the experiments that involved GDP-β-S, we effectively and selectively blocked all G-protein signaling that depends on Gα subunits, but only in the individual neuron that was patched.

The results of these experiments make two important points. First, we found that L-368 reproduced the effect of Oxtr-A when bath applied to PVN CRH neurons voltage clamped at −70 mV in slices extracted from animals that received hypertonic saline. Specifically, we noted a shift in current density of −1.8 ± 0.6 pA/pF (Fig. 4A-B, grey symbols). This effect is significantly bigger than 0 as determined by a one-sample t-test (n=5, p=0.04), and also is not significantly different than the response produced by bath application of Oxtr-A presented in Fig. 1D (−1.6 ±.7 pA/pF, n=8, p=0.85, two-sample unpaired t-test). Second, we found that this effect of L-368 was completely eliminated in cells that were patched with an internal solution that contained GDP-β-S (Δcurrent density = 0.4 ± 0.2 pA/pF, p=0.04, two-sample unpaired t-test vs. control condition, and p=0.08, one-sample t-test vs. null hypothesis of mean = 0). Collectively, these data reinforce the conclusion that tonic inhibition of PVN CRH neurons after acute salt loading depends on activation of Oxtr(s), and further indicate that the mechanism requires G-protein activation specifically in the patched neuron.

**Figure 4.**
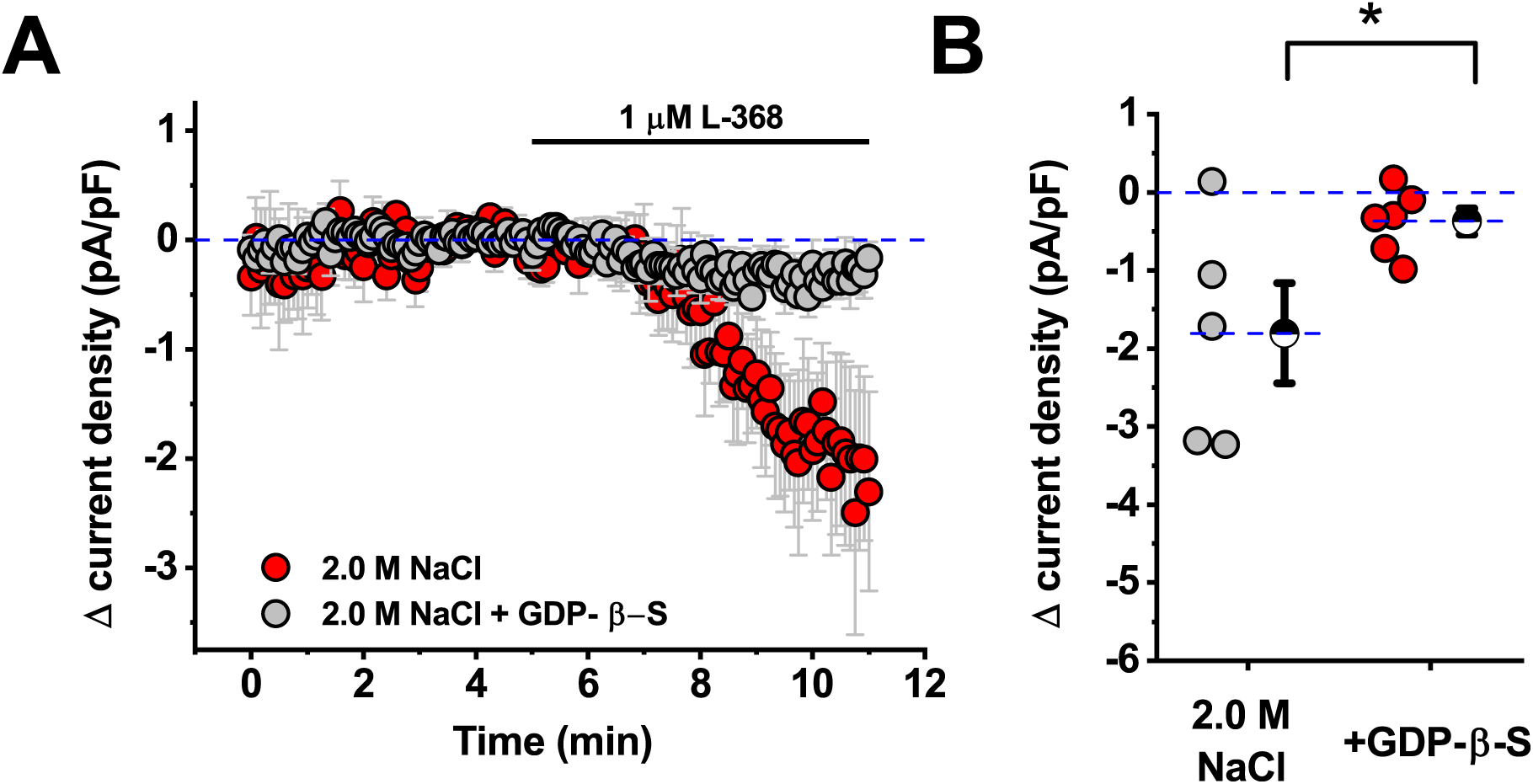
Inhibitory tone observed in CRH neurons from salt loaded animals is blocked by intracellular inhibition of GPCRs. **A)** Oxytocin receptor mediated inhibitory tone in PVN CRH neurons extracted from animals that received hypertonic saline (red symbols) is abolished in cells filled via the patch pipette with GDP-β-S (grey symbols). This result suggests that effect of Oxtr-A depends on GPCRs expressed by the patched neuron. **B)** Box plot illustrating specific change in current density produced by Oxtr-A for each cell that contributed to the summary plot in panel A. Large hatched symbols to the right of the data represent the mean of each group, and error bars indicate the SE. The asterisk indicates that GDP-β-S significantly reduced the effect of Oxtr-A (p=0.04). See results for additional analysis.

### Oxtr mRNA is expressed in the majority of PVN CRH neurons, and selective deletion of Oxtr(s) from CRH neurons reduces tonic inhibition produced by acute salt loading

While results presented above provide a compelling argument that Oxtr-mediated inhibition observed after acute salt loading requires direct activation of Oxtr(s) expressed by PVN CRH neurons, we performed two additional experiments to further validate that hypothesis. First, we used RNAscope in situ hybridization to determine whether PVN CRH neurons synthesize Oxtr mRNA (see Methods). We found co-localization of Oxtr mRNA in 82.6 ± 4.5% of the PVN CRH neurons that were sampled (Fig. 5A). Therefore, we next sought to test the hypothesis that selective genetic deletion of Oxtr(s) from CRH neurons would reduce or eliminate Oxtr mediated inhibition observed after acute salt loading. Towards that end, we crossed mice homozygous for Oxtr flox gene with mice heterozygous for the CRH-Cre gene to generate both mice lacking the Oxtr in CRH neurons (CRH-Oxtr KO mice), and the appropriate littermate controls harboring only the Oxtr-flox gene (see Methods for additional details). This breeding scheme created both mice that lack Oxtr in CRH neurons (CRH-Oxtr KO mice) and appropriate littermate controls (see Methods for additional details). Both CRH-Oxtr KO mice and littermate controls were rendered hypertonic following 2.0 M NaCl injection and putative CRH neurosecretory neurons were identified in vitro and tested with an oxytocin receptor antagonist as in earlier experiments. We found that PVN CRH-like neurons from littermate controls (n=8) did exhibit an Oxtr-dependent tonic inhibition that was blocked by L-368 (Δ current density = −0.91±0.37 pA/pF; one-sample t-test, p=0.04 vs. null hypothesis mean=0; Fig 5B-C). Further, this statistically significant effect was not present in PVN CRH neurons (n=6) from CRH-Oxtr KO mice (Δ current density = −0.23±0.14 pA/pF; one-sample t-test, p=0.16 vs. baseline). These data effectively indicate that selective deletion of Oxtr(s) from PVN CRH neurons prevents expression of significant Oxtr-mediated tonic inhibition after acute salt loading. Nevertheless, it should also be noted that there was not a significant difference between group means for CRH-Oxtr KO mice and littermates when compared directly to one another using a two-sample t-test (p=0.12).

**Figure 5.**
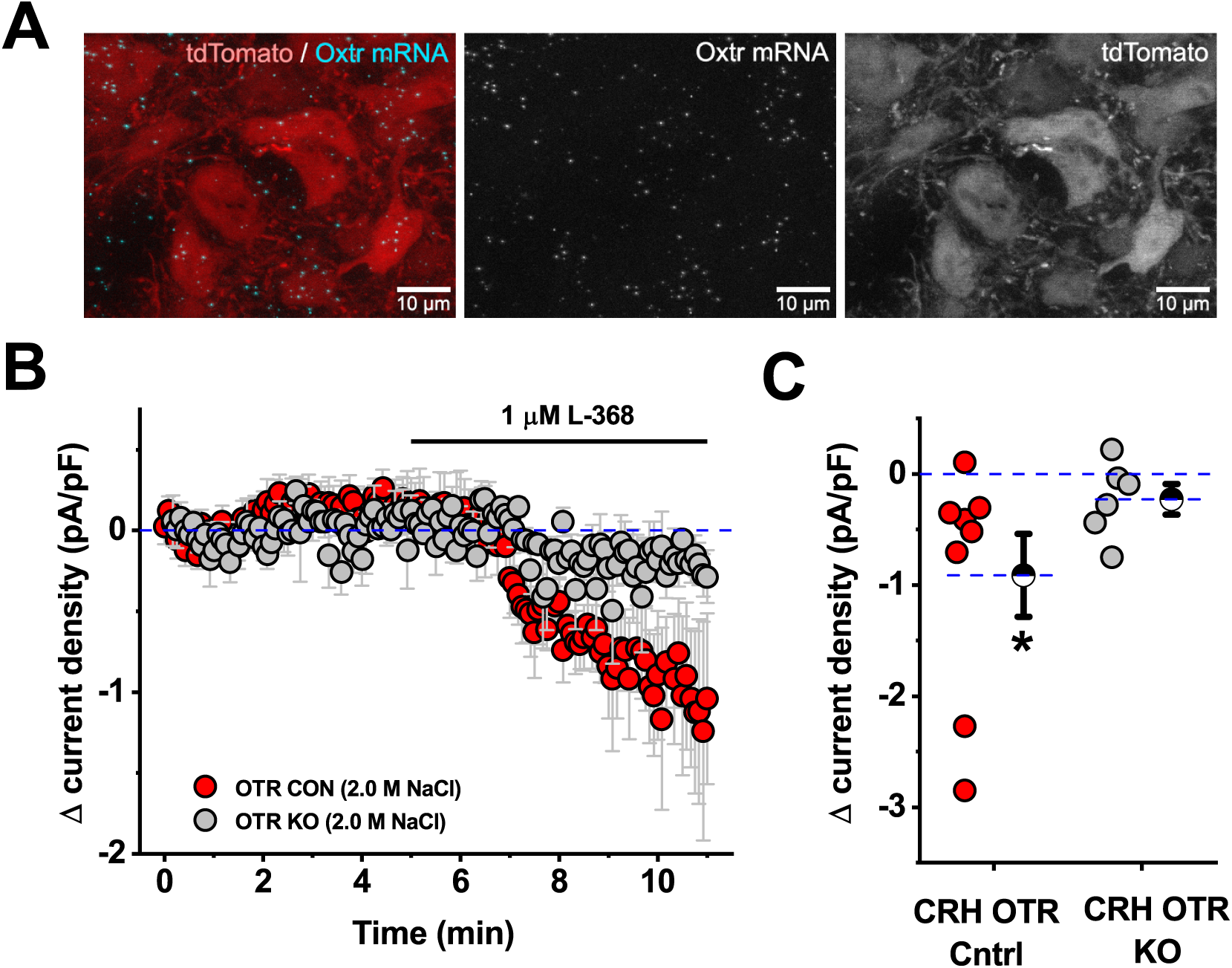
mRNA for Oxtr(s) is present in the majority of PVN CRH neurons, and selective deletion of Oxtr(s) from CRH neurons reduces tonic inhibition produced by acute salt loading. **A)** Fluorescent *in-situ* hybridization reveals Oxtr mRNA (cyan) in 82.6 ± 4.5% tdTomato-expressing (putatively CRH) neurons examined (red, 809 cells from 3 animals examined in total). Scale bar represents 10 µm. **B)** Conditional knockout of OTR in CRH neurons attenuates oxytocinergic tone in salt-treated animals compared to littermate controls. **C)** Box plot illustrating specific change in current density produced by Oxtr-A for each cell that contributed to the summary plot in panel D. Large hatched symbols to the right of the data represent the mean of each group, and error bars indicate the SE. The asterisk indicates that antagonism of Oxtr(s) still causes a significant reduction in current density in littermate controls (p=0.04) that was absent in CRH neurons from Oxtr-CRH KO animals. See results for additional details.

### Salt loading reduces excitability of PVN CRH neurons through opening an inwardly rectifying potassium channel

Prior work from our group has indicated that acute salt loading robustly inhibits activation of the HPA axis, as indicated by lower levels of ACTH and corticosterone subsequent to psychological stress (Krause et al., 2011; Frazier et al., 2013; Smith et al., 2014). If this effect is indeed produced by Oxtr mediated inhibition of PVN CRH neurons that initiate activation of the HPA axis, then it is desirable to understand in greater detail exactly how activation of Oxtr(s) on CRH neurons produces functional inhibition. As a first step towards that end, we carefully evaluated input resistance in PVN CRH neurons from animals that received isotonic vs. hypertonic saline. We found that acute salt loading significantly reduced whole-cell input resistance as measured from a brief voltage step from −70 mV to −80 mV (Isotonic Rm: 1225 ± 263 MOhm, n=7, Hypertonic Rm: 692 ± 58 MOhm, n=9, p=0.04). This finding suggests that salt loading opens a membrane channel that drives membrane potential more negative. Based on that observation, we next performed an additional series of voltage clamp experiments to evaluate the relationship between holding current and command potential between −110 mV and −50 mV (Fig. 6A). Although a two-way repeated measures ANOVA on data presented in Fig. 6A reveals no main effect of salt loading (F_1,14_=3.669, p=0.08), there was a significant salt × voltage interaction (F_1,9_=11.524, p<0.005). Post hoc tests revealed that PVN CRH neurons from salt loaded animals required significantly more current to hold the membrane potential at all voltages tested negative of −90 mV (p-values from 0.04 to << 0.01). Consistent with this observation, the difference between these two I-V plots, illustrated in Fig. 6B, further highlights that salt loading is associated with increased activation of an inwardly rectifying current that reverses near the equilibrium potential for potassium.

**Figure 6.**
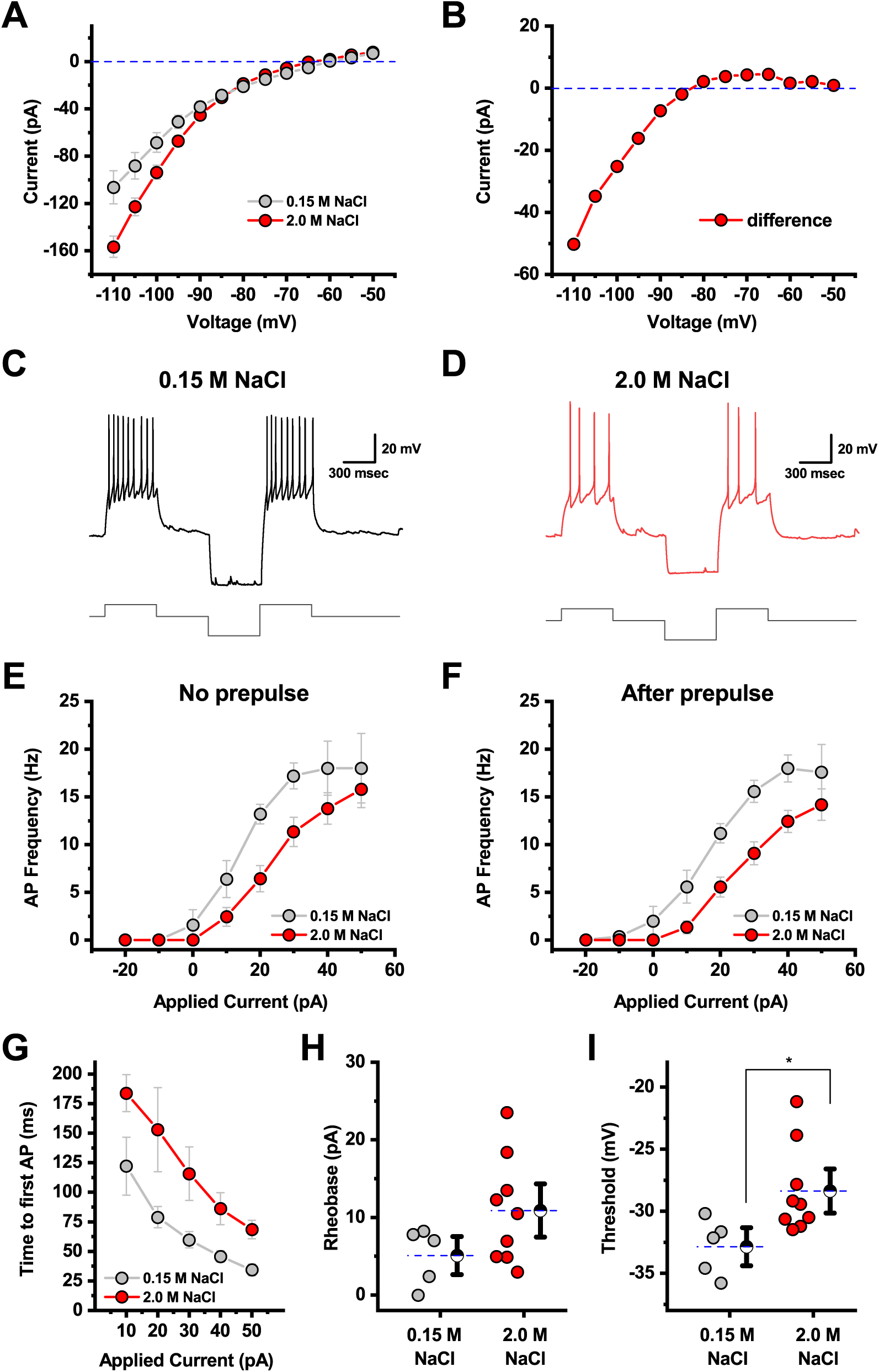
Salt loading reduces excitability of PVN CRH neurons through opening an inwardly rectifying potassium channel. **A)** Current/voltage relationship of PVN CRH neurons indicates more inhibitory current is required to hold neurons of salt-loaded animals at negative potentials. **B)** The difference of the current/voltage curves in A reveals salt loading activates an inwardly-rectifying current which reverses near the equilibrium potential for potassium. **C)** A representative current clamp recording showing the response to a PVN CRH neuron extracted from an animal that received isotonic saline to a +20 pA current injection delivered either in isolation, or after a −50 pA hyperpolarizing pulse. **D)** Representative trace from an identical experiment as panel C, in a PVN CRH neuron extracted from a salt loaded animal, highlights decreased responsiveness to a 20 pA current injection, without or with the hyperpolarizing prepulse. **E-F)** Complete neuronal gain curves constructed from cells recorded as in panels C-D indicate that salt loading produces a statistically significant reduction in neuronal gain. See results section for additional details. **G)** Acute salt loading increases time to first action potential in response to injection of 10 - 50 pA current steps. **H)** Salt loading increases action potential rheobase and threshold, as observed during a slow current ramp, although only the later effect was statistically significant (n=5, 9, p=0.11 and 0.03, respectively, two-sample unpaired t-test).

G-protein coupled inwardly rectifying potassium channels (GIRKs) often reduce neuronal excitability by shifting the resting membrane potential more negative and/or by decreasing neuronal input resistance, however, the effects of these changes on neuronal activity in physiological conditions can be complex. In order to more directly evaluate the effect of salt loading on excitability of PVN CRH neurons, we evaluated the response of the neurons to direct current injection in current clamp, with or without a hyperpolarizing prepulse of −50 pA. Fig. 6C illustrates the response of a representative PVN CRH neuron to a +20 pA current injection, both with and without the hyperpolarizing prepulse, in an animal that received isotonic saline. Fig. 6D illustrates reduced firing in response to an identical stimulus in a representative trace recorded from a PVN CRH neuron extracted from a salt loaded animal. Complete neuronal gain curves for isotonic and hypertonic groups with and without a hyperpolarizing prepulse indicate that there was a main effect of salt loading on action potential frequency (salt loading reduced neuronal gain, both without, and with, a hyperpolarizing prepulse, F_1,12_=7.99, p=0.02, and F_1,12_=12.86, p<0.005, respectively, Fig. 6E-F). We also noted that salt loading increased the time to first action potential, for current injections between 10 and 50 pA, with the difference being larger in response to smaller current injections (main effect of salt loading, F_1,6_=9.33, p=0.02, Fig. 6G). Similarly, we measured both the rheobase and the threshold of the first action potential observed during a slow current ramp (10 pA/sec)). Salt loading increased the rheobase from 5.1 ± 1.6 pA to 10.9 ± 2.3 pA, and increased the threshold from −32.9 ± 1.0 mV to −28.4 ± 1.2 mV, although only the later effect was statistically significant (n=5, 9, p=0.11 and 0.03, respectively). Collectively, these data further reinforce the conclusion that salt loading produces a clear and functional reduction in the excitability of PVN CRH neurons.

## Discussion

In this study we used a previously characterized CRH reporter animal (Cusulin et al., 2013; Smith et al., 2014; Chen et al., 2015; Walker et al., 2018) to provide the first direct demonstration that acute hypernatremia produces an Oxtr dependent inhibitory tone on identified parvocellular PVN CRH neurons. A first basic question to ask in considering this study is how fundamental information on the identity and physiological properties of PVN CRH neurons observed here aligns with prior data and general expectations. In that regard, it is worth highlighting that we found the vast majority of genetically identified PVN CRH neurons to be parvocellular neurons, with low whole-cell capacitance and high input resistance, that lacked both a clear A-type potassium current and a robust low threshold spike. In those respects, our findings are consistent with early characterizations of ‘Type II’ parvocellular neurosecretory neurons postulated to regulate HPA axis activation (Luther et al., 2002). Importantly, these results also align quite well with initial physiological characterizations of PVN CRH neurons in the identical CRH-ires-Cre mouse model as reported by other investigators (Cusulin et al., 2013). Notably, we also report that PVN CRH neurons rarely express NP1, while Cusulin et al (2013) reports PVN CRH neurons are robustly labeled by peripherally delivered fluorogold, and that they are rarely immunoreactive for either vasopressin or oxytocin. While it is possible that a small subset of PVN magnocellular neurons could express CRH, collectively, we believe the data described above make a compelling case that PVN CRH neurons as identified in this study are overwhelmingly parvocellular neurosecretory neurons. As such, we conclude that the mechanism revealed here very likely underlies the previously reported ability of acute hypernatremia to dampen activation of the HPA axis as produced by subsequent restraint stress (Krause et al., 2011; Frazier et al., 2013; Smith et al., 2014).

An important aspect of that mechanism, as described in the results section, is the idea that it depends on direct activation of Oxtr(s) that are expressed on the PVN CRH neurons themselves. In our view, multiple lines of evidence presented here strongly support that conclusion. Most directly, whole-cell recordings reveal an effect of a bath applied Oxtr antagonist on identified PVN CRH neurons extracted from salt loaded animals. This effect is largely insensitive to bath applied antagonists for glutamate and GABA receptors, does not require action potential dependent transmission in the slice, and is blocked by GDP-β-S delivered intracellularly just to the patched neuron. Further, in situ hybridization studies reveal clear expression of Oxtr mRNA in a majority of PVN CRH neurons examined, and conditional knockout of Oxtr from CRH neurons eliminates the effects of hypernatremia observed in vitro.

While we believe the data above provide a compelling argument that parvocellular PVN CRH neurons do indeed express functional Oxtr(s) that can be activated by acute hypernatremia, it is important to carefully consider two studies that at face value provide some potentially contradictory data. The first is a single cell RT-PCR study that reports that parvocellular PVN CRH neurons are Oxtr negative, but that there also exists a smaller population of CRH expressing mostly oxytocinergic magnocellular neurons that are Oxtr positive (Dabrowska et al., 2013). This study follows a prior report from the same authors indicating minimal somatodendritic overlap between PVN neurons immunoreactive for oxytocin vs. those immunoreactive for CRH, and that used single cell RT-PCR to reveal that 2 of 3 CRH positive PVN neurons examined (not explicitly defined as parvocellular vs. magnocellular) expressed mRNA for Oxtr(s) (Dabrowska et al., 2011). Both of those findings are highly consistent with present results. Later evidence suggesting that a small subset of PVN CRH neurons are in fact Oxtr-expressing magnocellular oxytocinergic neurons is also compatible with prior and current data, at least to the extent that these cells represent a very small minority of total PVN CRH neurons. However, the complete lack of Oxtr expression in presumed parvocellular PVN CRH neurons reported by Dabrowska et al 2013 is more difficult to reconcile with present data. As it stands, PVN neurons used for transcriptomic analysis in Dabrowska et al. (2013) were initially defined as magnocellular vs. parvocellular based purely on somatic size and spatial location within the PVN, which might lead to somewhat different cell selection than in other studies. On the other hand, electrophysiological properties reported for parvocellular CRH neurons were generally consistent with expectations. Overall, it seems possible that very low levels of Oxtr expression in PVN CRH neurons could make reliable detection with either single cell RT-PCR or in situ hybridization extremely challenging, or that there is some stress related plasticity in Oxtr expression in PVN CRH neurons that is not fully revealed in the present study.

A second highly relevant study using largely in vitro electrophysiological techniques similarly reports that PVN CRH neurons are Oxtr negative based on the observation that bath applied oxytocin has minimal if any effect on intrinsic excitability of these neurons (Jamieson et al., 2017). The most notable difference between the approach of Jamieson et al (2017) and our own work is that Jamieson et al (2017) relied on bath application of oxytocin (1 μM for 5 minutes) to activate putative Oxtr(s), while we relied on acute hypernatremia to promote natural release of endogenous oxytocin. Indeed, we acknowledge that in pilot studies, we have also been unable to directly or reliably mimic the effects of endogenous oxytocin on PVN CRH neurons with a bath applied exogenous Oxtr agonist. However, there are several complexities with this approach. Notably, exogenous peptides stick to tubing and glassware, are sensitive to temperature, and are broken down by endogenous peptidases. To compensate for these types of issues, experiments with bath applied peptides commonly overshoot effective concentrations of endogenous agonists, yet this may lead to additional problems because Oxtr(s) are likely subject to rapid internalization and/or desensitization in response to high concentrations of agonist (Raggenbass et al., 1998; Robinson et al., 2003; Smith et al., 2005; Terenzi and Ingram, 2005). Indeed, despite broad expression of Oxtr(s) in brain, there remain relatively few studies that show robust responses to bath application of exogenous Oxtr agonists using whole-cell recordings in vitro. Further, many that do reveal transient and/or non-repeatable responses to lower concentrations than those employed by Jamieson et al (2017), and almost all involve activation of Oxtr(s) coupled to effectors that promote depolarization rather than hyperpolarization. Therefore, in our view, the difficulty involved in modeling endogenous oxytocin release in the PVN as produced by acute hypernatremia with a bath applied exogenous agonist does not substantially or directly reduce the value or power of multiple other lines of evidence described above, which do point towards functional Oxtr(s), capable of responding to endogenous oxytocin release, being expressed by parvocellular PVN CRH neurons.

Next, it is important to consider how data presented here aligns with prior studies that have directly implicated central Oxtr(s) in modulation of stress responsiveness. As noted in the introduction, significant prior evidence indicates that a variety of acute psychological stressors promote central oxytocin release (Ježová et al., 1993; Nishioka et al., 1998; Wotjak et al., 1998; Engelmann et al., 1999; Torner et al., 2017), and substantial additional data indicates that centrally acting oxytocin can modulate stress induced activation of the HPA axis (Windle et al., 1997, 2004; Neumann et al., 2000). In most prior studies, a specific mechanism for Oxtr dependent modulation of stress induced activation of the HPA axis is not directly revealed, although several lines of evidence have promoted the idea that, either directly or indirectly, decreased synthesis of CRH in PVN CRH neurons is involved (Nomura et al., 2003; Jurek et al., 2015; Winter and Jurek, 2018). In considering our data in light of these studies it is important to emphasize that acute hypernatremia is a peripheral non-psychological stressor that is likely to have very different effects on PVN activity than the psychological stressors previously used to promote central oxytocin release. Indeed, we would hypothesize that hypernatremia is likely to evoke central oxytocin release either via direct osmotic regulation of PVN magnocellular oxytocinergic neurons, or via excitatory input from osmosensitive neurons in circumventricular organs. By contrast, we expect oxytocin release in the PVN following psychological stressors is likely to involve activation of substantially more descending central inputs into the PVN, and is also likely to be confounded with slower genomic effects ultimately produced by circulating glucocorticoids. That said, it seems possible that oxytocin acting locally in the PVN could promote Oxtr-mediated genomic modulation of CRH synthesis, and that this would ultimately act synergistically with mechanisms revealed here to reduce the HPA axis response to additional stressors. Nevertheless, in a broader sense, we believe the results presented here emphasize a new context for Oxtr mediated modulation of the HPA response to stress; one that is not initiated by prior psychological stressors, one that has potential to work on shorter time scales, and one that has some interesting potential social relevance (Smith et al., 2015).

Finally, it is worth considering what this study helps reveal about differences between acute and chronic hypernatremia with respect to modulation of the HPA axis. Prior studies indicate that chronic salt loading alters CRH expression patterns in hypothalamus (Young, 1986), is associated with reduced ACTH release from the anterior pituitary (Dohanics et al., 1990; Chowdrey et al., 1991), does not alter stress induced release of oxytocin despite increasing basal levels of oxytocin in plasma (Chowdrey et al., 1991), and yet does promote sensitization of the HPA axis during rehydration (Amaya et al., 2001). Recent work from our group has revealed that 5 days of chronic salt loading produces a moderate reduction in CRH synthesis in neurosecretory regions of the PVN, a clear reduction in sEPSC frequency recorded from PVN CRH neurons, and an attenuation in the HPA axis response to restraint stress that was apparent during recovery rather than immediately after restraint (Krause et al., 2017). Broadly speaking, these results reinforce the idea that acute Oxtr mediated modulation of PVN CRH neurons as detailed here is a mechanism likely associated with acute paracrine release of oxytocin in the PVN, and that distinct mechanisms underly longer term adaptation to chronic salt and/or or subsequent rehydration.

## Acknowledgements

This study was supported by AHA predoctoral fellowship 13PRE17100047 (DP), NIH grants MH104641 (CJF), HL096830 (EGK), HL122494 (EGK), and K99/R00-HL-125805 (ADdK).

## Conflict of interest

The authors declare no conflicts of interest.

